# Inferring weighted gene annotations from expression data

**DOI:** 10.1101/096677

**Authors:** Michael Cary, Cynthia Kenyon

## Abstract

Annotating genes with information describing their role in the cell is a fundamental goal in biology, and essential for interpreting data-rich assays such as microarray analysis and RNA-Seq. Gene annotation takes many forms, from Gene Ontology (GO) terms, to tissues or cell types of significant expression, to putative regulatory factors and DNA sequences. Almost invariably in gene databases, annotations are connected to genes by a Boolean relationship, e.g., a GO term either *is* or *isn’t* associated with a particular gene. While useful for many purposes, Boolean-type annotations fail to capture the varying degrees by which some annotations describe their associated genes and give no indication of the relevance of annotations to cellular logistical activities such as gene expression. We hypothesized that weighted annotations could prove useful for understanding gene function and for interpreting gene expression data, and developed a method to generate these from Boolean annotations and a large compendium of gene expression data. The method uses an independent component analysis-based approach to find gene modules in the compendium, and then assigns gene-specific weights to annotations proportional to the degree to which they are shared among members of the module, with the reasoning that the more an annotation is shared by genes in a module, the more likely it is to be relevant to their function and, therefore, the higher it should be weighted. In this paper, we show that analysis of expression data with module-weighted annotations appears to be more resistant to the confounding effect of gene-gene correlations than non-weighted annotation enrichment analysis, and show several examples in which module-weighted annotations provide biological insights not revealed by Boolean annotations. We also show that application of the method to a simple form of genetic regulatory annotation, namely, the presence or absence of putative regulatory words (oligonucleotides) in gene promoters, leads to module-weighted words that closely match known regulatory sequences, and that these can be used to quickly determine key regulatory sequences in differential expression data.

## Introduction

Genome annotation is a central task in modern biology, and a great deal of effort has been dedicated to the construction of controlled vocabularies to describe cellular organization and function, and to the association of terms in these controlled vocabularies to specific genes. Because annotating genes empirically is costly and time consuming, computational methods have been developed to infer gene annotations from existing annotations and additional data, such as gene sequences, interaction network connectivity, and gene-expression profiles^1–5^. To date, the aim of such methods has been to decide whether a gene should or should not be annotated with a particular term. While some of these methods provide a confidence score for their predictions, such a score does not necessarily reflect the relevance of the annotation to the gene’s role in the cell. For example, an enzyme could confidently be predicted to have a particular catalytic activity through sequence analysis, but this activity could be vestigial and unrelated to the enzyme’s current role. Similarly, *in vitro* transcription factor binding assays, such as those generated by the ENCODE and modENCODE projects^6,7^, can be used to annotate genes with lists of transcription factors that bind proximally, but the relative importance of each factor to the regulation of each gene would not be clear from such data alone. We hypothesized that relevance scores for gene annotations could be useful both for understanding the functions of individual genes, and for interpreting data from genome-scale assays. One way to generate such scores would be to determine how often the cell behaves as if a particular annotation is an organizing principle of its activity. We conjectured that a large, high-quality set of gene expression modules could be used to make this determination and developed a method to generate weighted annotations using a compendium of gene expression data and a set of Boolean annotations.

Our method consists of three main steps (Fig. 1). In the first, we predict gene transcription modules (sets of co-regulated genes) using independent component analysis (ICA) of a large compendium of expression data. ICA has been applied to gene module prediction before^8–12^, but we have refined the process in a way that improves the results substantially according to several different measures. These predicted gene modules serve as an intermediate data structure in our algorithm for module-weighted annotations, but they are revealing in their own right, and provide new insights into properties of gene expression, some of which we present here.

**Figure 1.**
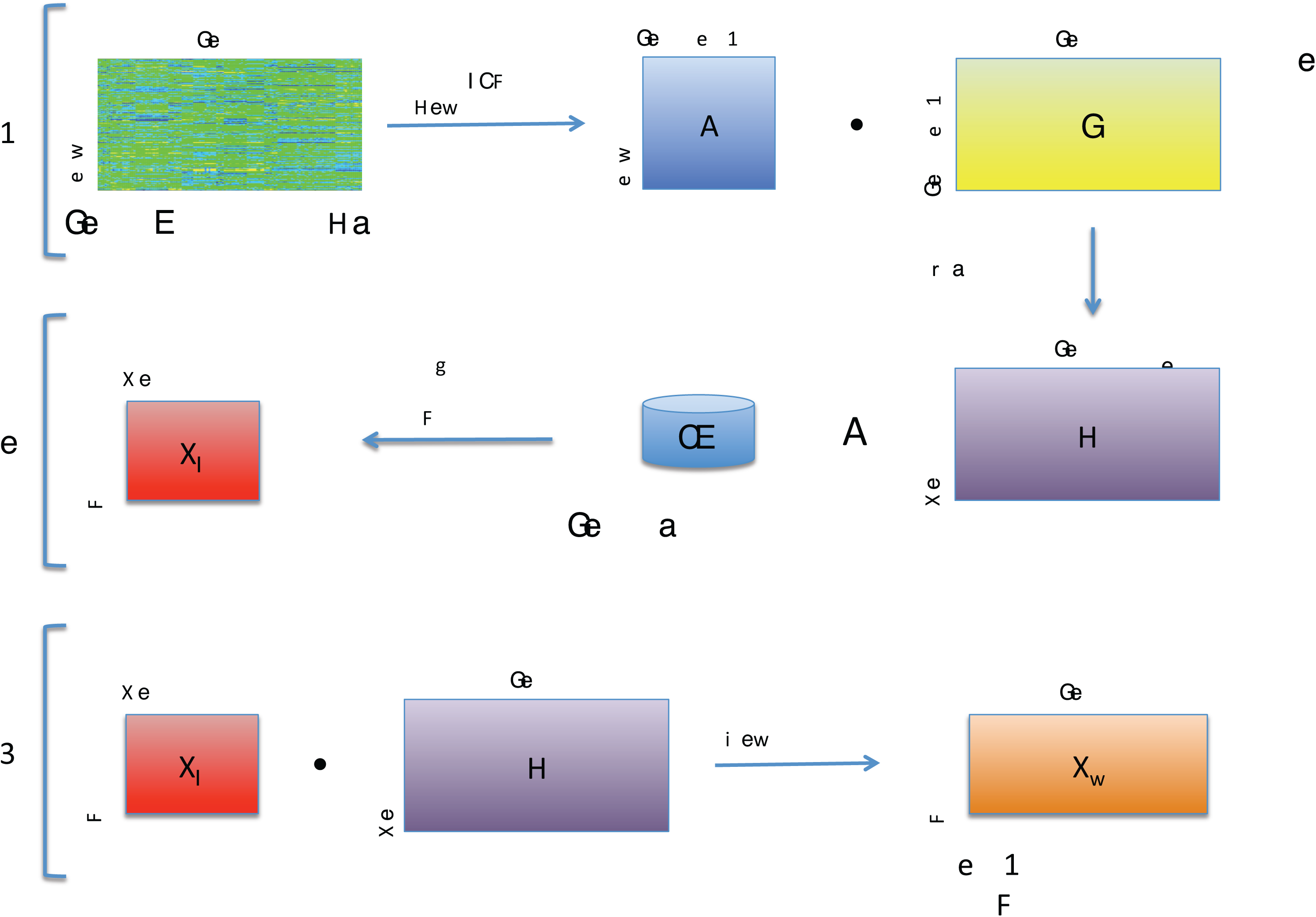
Figure 1.

In the second step, we calculate the enrichment or depletion of each annotation in each gene module using hypergeometric statistics, and construct a matrix with this data using the negative log of enrichment p-values (or the positive log of depletion p-values, if they are smaller). Finally, we determine the matrix product of the gene module definition matrix from step 1 and the annotation enrichment matrix resulting from step 2. This yields a matrix relating each annotation to each gene, i.e., module-weighted gene annotations.

We check the consistency of module-weighted annotations with existing knowledge by inspecting strongly positive and strongly negative genes for several different annotations, and inspecting which annotations are most strongly weighted for several well-characterized genes. We then test the utility of module-weighted annotations for gene set enrichment analysis (GSEA) by assessing their performance on a large set of gene expression experiments. We show that using module-weighted annotations for GSEA appears less prone to the confounding effect of gene-gene correlations than traditional GSEA.

Finally, we generate module-weighted annotations for a simple form of genetic regulatory annotation: the presence or absence of putative regulatory words (oligonucleotides) in gene promoters. We show that module-weighted words can reveal important regulatory sequences in gene expression data by using them to analyze data from several experiments, including *C. elegans isp-1* (respiratory chain) and *hif-1* (hypoxia-inducible transcription factor) mutant microarrays. Mutation of the *isp-1* gene extends lifespan in a *hif-1*-dependent fashion^13–15^, but the significant gene sets from microarray data for these two mutants have little in common. In contrast, our method reveals extensive gene co-regulation; and in addition, several words are predicted to exert strong effects in both data sets, and these words match the canonical HIF-1 binding site.

## Results

### Optimization of genetic regulatory module prediction

Our algorithm relies on accurate predictions of genetic regulatory modules. A large body of gene expression data is publicly available^16,17^ and has enabled computational prediction of gene modules (co-regulated genes) by several groups^8,18–30^. Preliminary experimentation with published methods led us to choose ICA for performing module prediction, as modules predicted with ICA yielded stronger oligonucleotide enrichment in promoter regions than did modules predicted with the other methods we tested (Fig. 2e and additional data not shown; see Lee & Batzoglou^12^ for additional comparisons of ICA to other methods).

**Figure 2.**
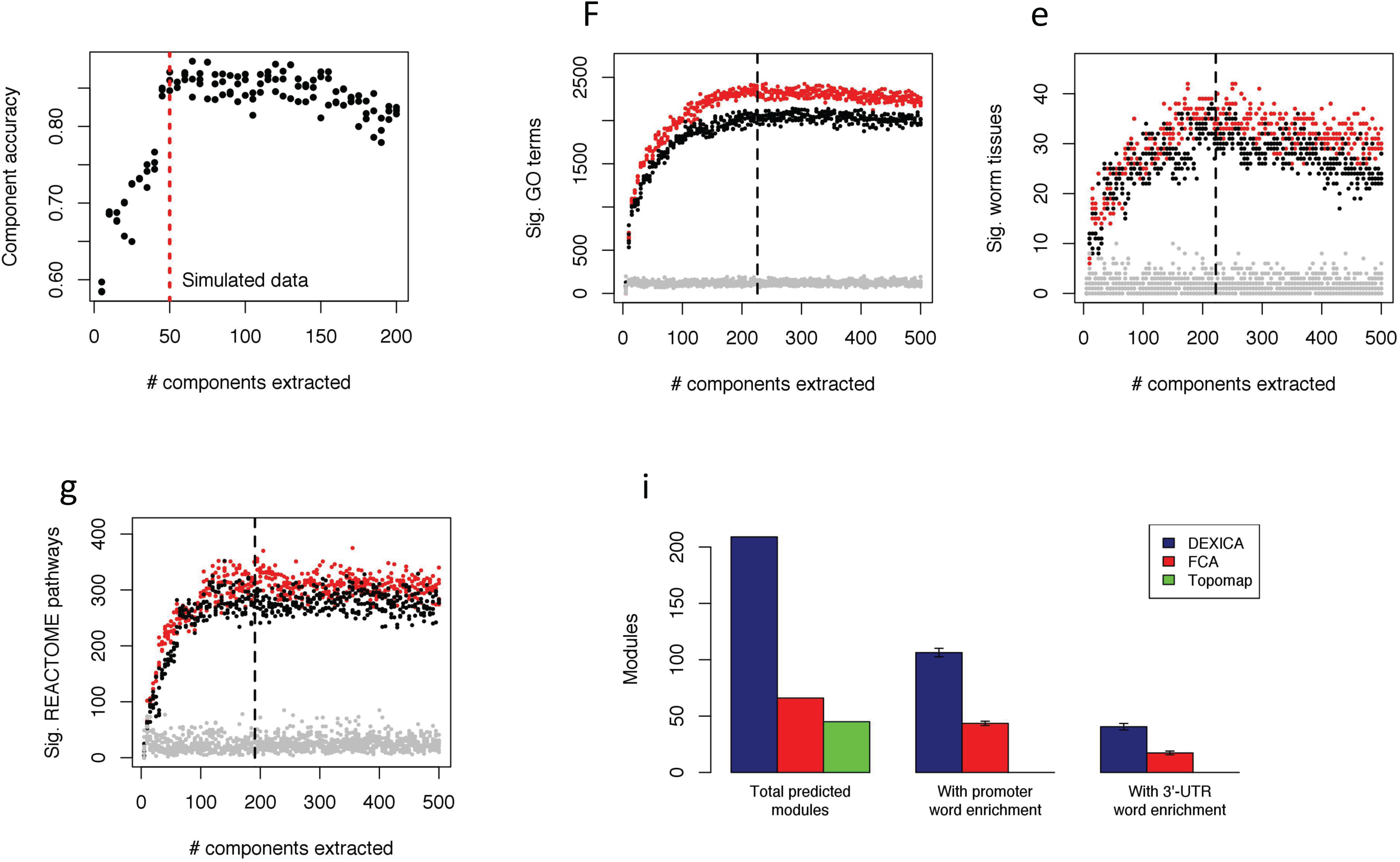
Figure 2.

Briefly, ICA is a blind source separation method that attempts to “unmix” a signal comprising additive subcomponents by separating it into statistically independent source signals^31,32^. In the common notation, a data matrix, ***X***, comprising multiple observations of a multidimensional variable, ***x***, is decomposed into two new matrices, a mixing matrix, ***A***, and a source matrix, ***S***:

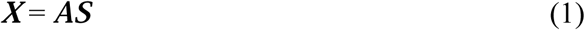

The ***A*** matrix contains the weight of each independent component in each observation, and the ***S*** matrix contains the weight of each element of ***x*** in each independent component. In the context of gene expression analysis, the elements of ***x*** correspond to genes, the observations correspond to genome-wide gene expression measurements, such as microarrays, and the independent components are interpreted as gene modules (essentially, genes whose expression levels change similarly across multiple arrays). The values in the ***S*** matrix correspond to the relative levels of inclusion of each gene in each gene module^10,33^.

Our preliminary investigation indicated that the performance of ICA was sensitive both to data preprocessing and to the number of components extracted. For example, using simulated data we found that module prediction accuracy was highest when the number of extracted components matched the true number of modules (Fig. 2a). Therefore, we first sought to optimize gene module prediction through ICA, evaluating results using biological information, including Gene Ontology (GO) term enrichment^34^, REACTOME pathway enrichment^35^, and tissue-specific expression enrichment in predicted gene modules. We applied our optimization strategy to a compendium of 1386 *C. elegans* Affymetrix arrays^36^, which we obtained from the Gene Expression Omnibus (GEO) database^17^. Our preliminary results indicated that applying dimension reduction procedures on the data matrix prior to performing ICA reduced the number of biologically significant components in the end result (data not shown), so we chose to optimize ICA of the full data matrix of 1386 arrays. We found that the highest quality modules were produced when we omitted between-experiment quantile normalization from the preprocessing procedure (see Methods) and when the number of extracted components (i.e., gene expression modules, or sets of co-regulated genes) ranged from 191 to 226 (Fig. 2b-d).

All of our module quality measures required translation of the independent components generated by ICA into discrete sets of genes, a process we refer to as partitioning. Typically, each component (gene module) is partitioned into three sets of genes: one set consisting of genes excluded from the module, and two other sets consisting of genes regulated in opposite directions. We refer to these latter two sets as “hemi-modules”, one set consisting of genes with highly positive weights and another consisting of genes with highly negative weights (assigned arbitrarily) in the independent component. Others have used a fixed threshold approach to partitioning^8,12,37^, for example, defining genes with weights exceeding +/− 3 standard deviations from the component mean to be “in” each hemi-module, and this is the approach we applied in figures 2a-c. However, we found that individual modules showed maximum annotation enrichment at different thresholds, suggesting that a ‘one-size-fits-all’ approach to partitioning is sub-optimal. An alternative approach to partitioning that we attempted (described in Frigyesi *et al.*^38^) failed to converge in many cases. Therefore, to increase partitioning accuracy, we trained an artificial neural network (ANN) to predict thresholds for partitioning of each component from the skewness and kurtosis of its weight distribution (see Supplemental Methods). Using this artificial neural network for partitioning in our optimization process produced similar results qualitatively (Fig. S1a-c), but resulted in a greater number of significant annotations across the range of parameters tested than did threshold-based partitioning (p < 2.2E-16, Fig. S1d). Therefore, we used ANN partitioning in the subsequent steps of our algorithm.

The mean optimum number of extracted components, determined using the quality measures we applied (dashed vertical lines in Fig. 1b-d and S1a-c), was similar for both threshold and ANN partitioning (209, and 209.33, respectively); therefore, we chose 209 as the final number of components to extract from the *C. elegans* Affymetrix microarray compendium. We refer to our process of extracting the optimal number of gene modules from a non-dimension-reduced gene-expression compendium using ICA as DEXICA, for <u>d</u>eep <u>ex</u>traction <u>i</u>ndependent <u>c</u>omponent <u>a</u>nalysis.

### Gene module validation

To test the prediction that the independent components generated by DEXICA correspond to genetic regulatory modules, we checked each module for enrichment of regulatory sequences in the promoter regions and 3’ untranslated regions (3’-UTRs) of module genes. To do this, we first generated a list of potential regulatory oligonucleotide sequences (called ‘words’) by applying the Mobydick algorithm^39^ to the set of all predicted ***C. elegans*** promoter regions, which we defined as the region extending from the transcription start site to 2000 base pairs upstream. (Many ***C. elegans*** regulatory sequences are located in this interval; however, we note that this method will exclude potential promoter sequences located exclusively upstream or downstream of this region.) We generated a second oligonucleotide list using the set of all predicted ***C. elegans*** 3’- UTRs (see Supplementary Methods). We then calculated the statistical significance of the over- or under-representation of genes bearing each word in each gene module (see Methods), using the hypergeometric test and the Simes method^40^ for multiple hypothesis testing (alpha level = 0.05), to determine the number of significant modules. Across multiple runs of DEXICA, the mean number of gene modules containing significant promoter words and 3’-UTR words was 106.3 and 40.6, respectively, which was significantly greater than that produced by other module prediction methods we tested (p < 2.2E-16, Fig. 2e).

Because the ICA algorithm that we employed during module prediction, *fastICA*, converges to a final solution from a random starting point^41^, small differences typically exist in the output of different runs of the algorithm; these differences can be seen in the vertical spread of data points in figures 2a-d, and in the error bars of figure 2e. While others have reconciled such differences through a clustering approach applied to the output of numerous runs of the algorithm (so called “iterated ICA”)^8,38^, when applied to our *C. elegans* Affymetrix compendium, we found that many of the final components generated by this method were highly correlated to one another, indicating non-independence and potential redundancy among the components (data not shown). We therefore sought to choose a single, high quality, *fastICA* run output to use as predicted gene modules. Because we considered word enrichment the most trustworthy measure of module quality, and because we observed a significant correlation (R = 0.27, p = 6.5E-3) between the total number of significant promoter words and the total number of significant 3’-UTR words in the results of different ICA runs with the same parameters (Fig. S2), as our final module set, we chose the run from a set of 100 with the best average rank in terms of significant promoter words and significant 3’-UTR words. This set ranked first in significant promoter words and third in significant 3’-UTR words.

### Global properties of gene expression revealed by predicted gene modules

Gene modules are sets of genes that are co-expressed. Unexpectedly, during our analysis of 3’-UTR word enrichment within gene modules, we observed that some modules appeared to be enriched for genes with long 3’-UTRs. To determine if this trend was statistically significant, we calculated the mean 3’-UTR length of each hemi-module and determined a p-value for length bias using Student’s t-test (Fig. 3a). Of the 418 hemi-modules, 65 contained a significant (q < 0.1) bias toward long 3’-UTR genes and 58 contained a bias toward short 3’-UTR genes.

**Figure 3.**
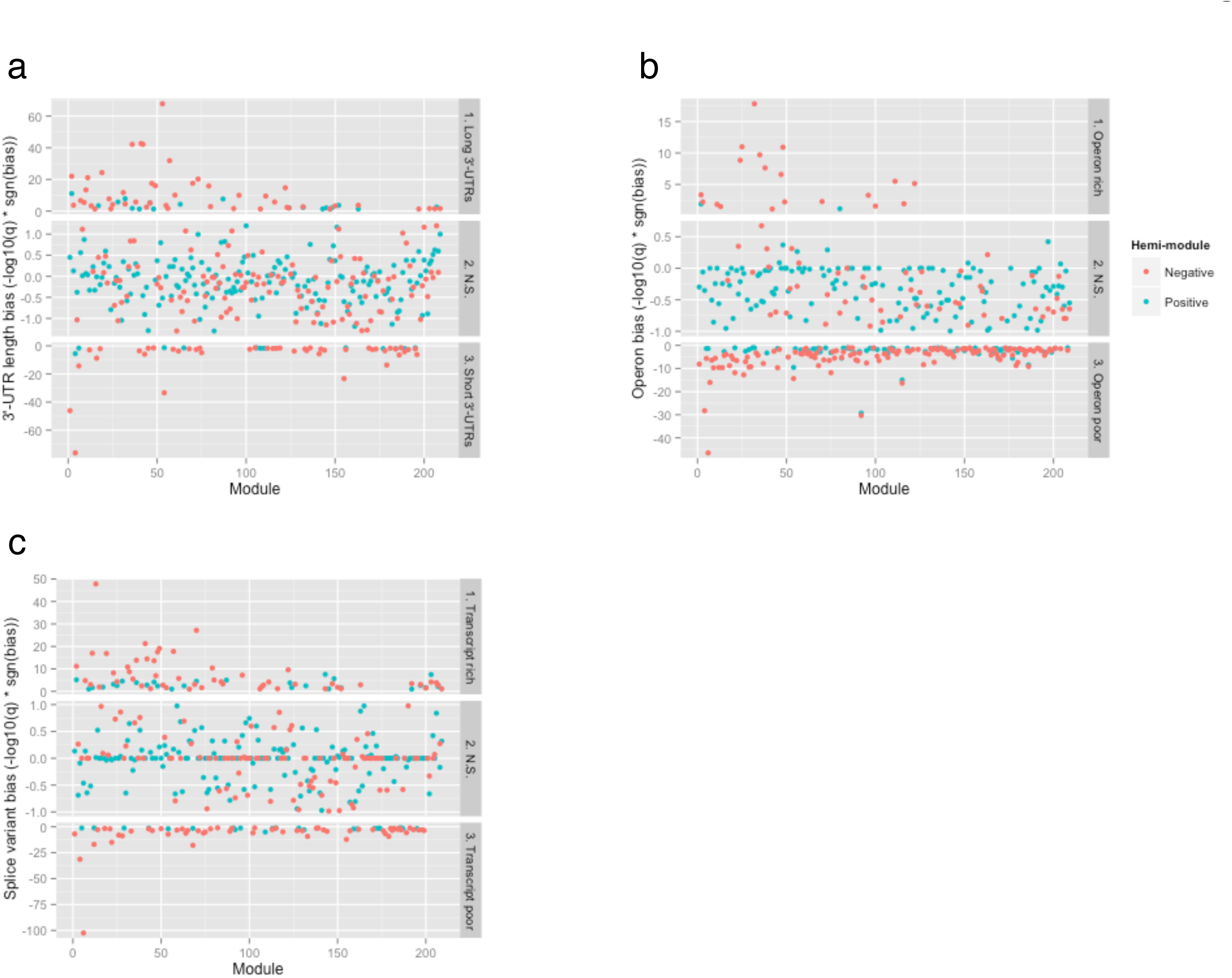
Figure 3.

To see if other gene-structure properties were enriched in specific gene modules, we tested each hemi-module for over- and under-enrichment of genes appearing in operons and for genes with multiple splice forms. Twenty-one hemi-modules were significantly enriched and 205 hemi-modules were significantly under-enriched for operon genes, and 81 hemi-modules were enriched and 80 hemi-modules were under-enriched for genes with multiple splice variants (Fig. 3b-c). Control tests performed on the same module set but with randomly scrambled gene IDs produced no significant modules for any of the gene properties we tested (Fig. S3). Taken together, these results suggest that genetic regulatory modules tend to comprise genes with gross similarities in gene structure. This association, in turn, raises the possibility that these shared structural features (long 3’-UTRs, etc.) house important biological information, either for gene regulation or gene function. Consistent with this idea, genes within operons are enriched in the set of ***C. elegans*** genes switched on during recovery from growth-arrested states^42^, and 3’-UTR length has been shown to be anti-correlated with mRNA stability, as longer 3’-UTRs are more subject to micro-RNA mediated repression than are shorter 3’-UTRs^43^.

### Generation of module-weighted annotations

Our calculation of module-weighted annotations takes advantage of the fact that, in modules generated by ICA, each gene has a weight in each module. Given a score or weight for each annotation in each module, this allows genes to be associated with annotations via a matrix product calculation. In our method, we create a matrix, ***X_a_***, comprising enrichment scores for each annotation / hemi-module combination (see Methods). Values in this matrix are derived from the log of enrichment p-values; highly positive values correspond to strongly enriched annotations in a hemi-module, highly negative values correspond to strongly under-enriched annotations. We transform the gene module matrix, ***S***, into a hemi-module matrix, ***H***, by concatenating it with a negative copy of itself row-wise:

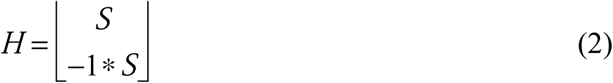

The product of this matrix with the ***X_a_*** matrix produces matrix ***X_w_***, which relates genes to annotations:

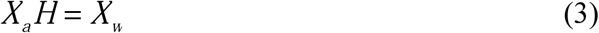

As a final step, we normalize the values in this matrix column-wise, i.e, separately for each annotation, by subtracting the mean and dividing by the standard deviation.

### Examination of module-weighted annotations

As an initial test of our method, we calculated module-weighted annotations for GO terms. We restricted the GO term set to those with at least 15 annotated genes in order to focus the analysis on robust signals; this set comprised 1651 GO terms. We first examined whether module-weighted annotations recapitulated Boolean annotations by testing whether genes associated with each term tend to have larger module-weighted annotations than other genes (two-sample KS test, alpha level = 0.05); this was true in 98.6% (1628/1651) of cases.

We then tested which GO terms contained the most strongly-weighted genes when normalization was omitted from their calculation, with the expectation that these should correspond to processes that are tightly regulated at the level of gene transcription. The GO terms with the most highly weighted genes were mainly associated with ribosome and/or nucleosome structure and function, and the most highly weighted genes for these terms were known ribosomal or nucleosome genes (Table 1).

We next examined which GO terms did not have any strongly weighted genes. Many of these terms were associated with signal transduction and kinase activity (Table 1). To test whether these observations were statistically significant, we ranked each GO term by the weight of its most strongly associated gene. We then constructed a list comprising all words used in all GO terms, excluding words shorter than three letters and uninformative words (e.g., “the” and “for”.) Finally, we tested each word for bias toward appearing near the top or bottom of the ranked GO term list. In agreement with our initial observations, the most significantly top-biased words pertained to macromolecular complexes, such as “nucleosome”, “cilium”, and “ribosomal”, and the most significant bottom-biased words pertained to cell signaling, such as “signal”, “kinase”, and “receptor” Many of the GO terms containing cell signaling words were generic in nature, e.g., *protein kinase regulator activity,* thus, our results may partially be explained by a lack of co-regulation among constituents of different signaling pathways. However, some specific cell signaling terms, e.g., *Notch signaling pathway* also appeared near the bottom of the ranked GO term list, suggesting that the genes annotated with such terms are either not strongly co-regulated at the gene expression level or that the biological conditions represented by the compendium did not perturb their expression enough to form modules with our method.

### Application of module-weighted annotations to gene set enrichment analysis

To test our hypothesis that module-weighted annotations could aid the interpretation of gene expression data, we devised an analysis method that makes use of weighted-annotations (see Methods) and compared its performance to traditional gene set enrichment analysis (GSEA)^44^. Our performance comparison is based on the assumption that pairs of highly similar experiments should show a strong correlation in the significance level of annotations (e.g., GO terms), and that highly dissimilar experiments should show no such correlation.

From a large set of gene expression fold changes, we selected the 100 most similar experiments (Spearman correlation, ρ, of gene fold changes near +/− 1) and the 100 least similar experiments (ρ near 0). We found that highly similar experiments showed a stronger correlation in the significance level (absolute value of z-scores) of annotations when using weighted-annotation analysis than when using GSEA (p = 9.5E-6). Likewise, we found that highly dissimilar experiments showed a weaker correlation in the significance level of annotations using our method (p < 2.2E-16, Fig. 4a). Indeed, the mean correlation between absolute values of z-scores for highly dissimilar experiments was close to zero for our method (mean ρ = 0.019), but significantly higher than zero for GSEA (mean ρ = 0.196). These results suggest that module-weighted annotations may provide resilience to the confounding effect of highly coordinated expression among genes annotated with certain GO terms, a known issue with GSEA (see Discussion), and that analysis of gene expression data using module-weighted annotations may provide more reliable biological insights than traditional gene set enrichment analysis.

**Figure 4.**
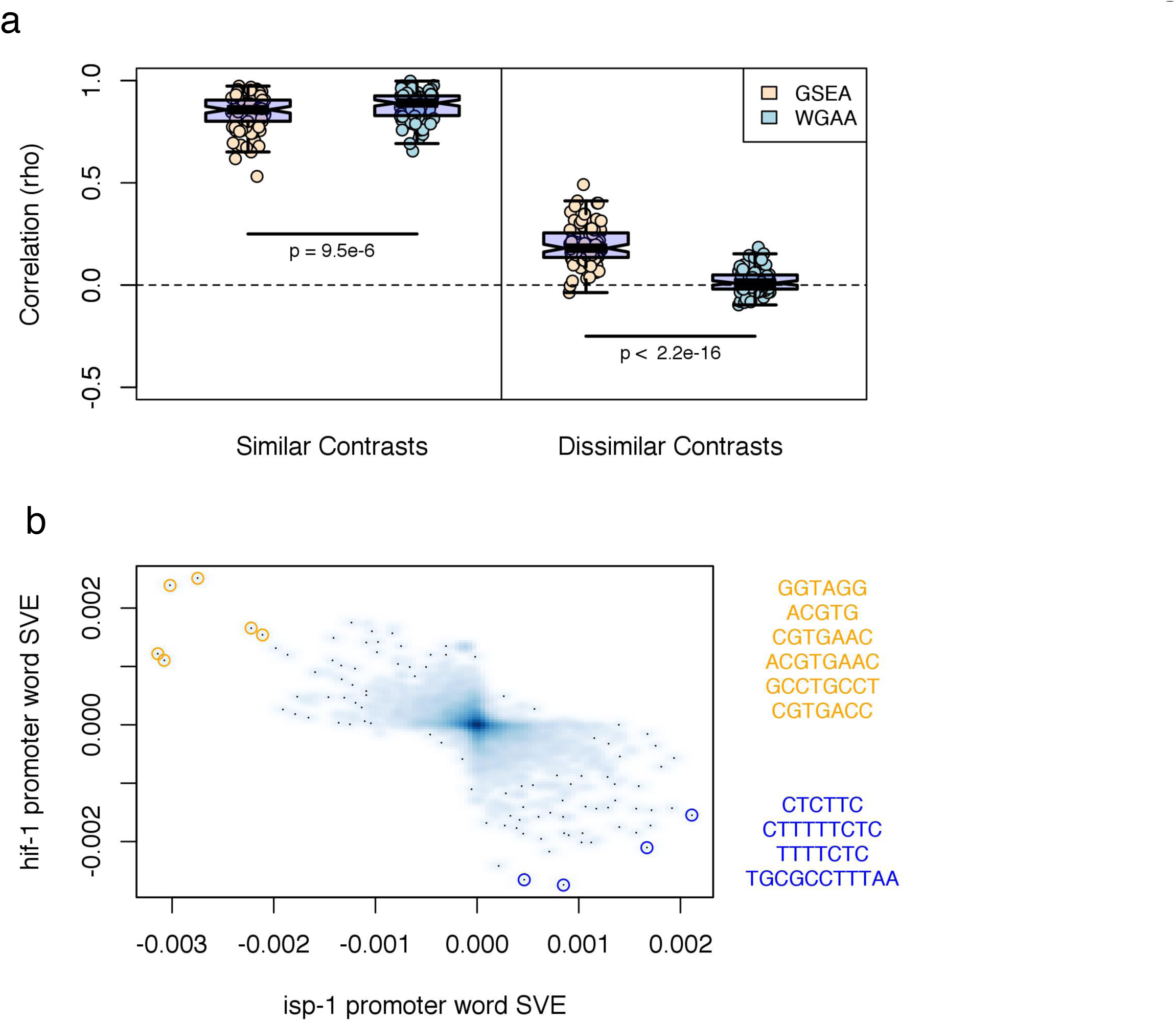
Figure 4.

### Module-weighted regulatory sequences

While any Boolean gene annotation may be converted into a module-weighted annotation with our method, it occurred to us that module-weighted regulatory sequences (i.e., a numeric indication of the degree to which a particular regulatory sequence appears to have an influence on a gene’s expression, given a set of gene modules), might be particularly useful for gene-expression analysis. To test this, we generated weighted annotations for each of the 5230 words in the promoter word dictionary (described above).

To validate predicted regulatory word weights, we searched the literature, the JASPAR database of transcription factor binding profiles^45^, and the GEO database for *C. elegans* transcription factors with both an experimentally-characterized DNA-binding profile and, separately, a microarray experiment that measured gene expression in a loss-of-function mutant of the transcription factor gene. This search yielded 6 transcription factors: *daf-12* (abnormal dauer formation), *daf-16/FoxO* (stress response and longevity), *hif-1* (hypoxia-inducible transcription factor)^46^, *hlh-30* (helix loop helix), *lin-14* (abnormal cell lineage), and *nhr-23* (nuclear hormone receptor). We then calculated two z-scores for each word in each loss-of-function microarray assay using the method described above: one score for the up-regulated genes, and another for the down-regulated genes (see Methods). Analyzing positively- and negatively-changing genes separately avoids a possible “cancelling out” effect that could arise when a word is enriched in both the positively and the negatively changing genes in a set of gene fold-changes. We then compared the top-scoring words to the DNA-binding profiles of the respective transcription factors. If the predicted promoter-word weights are accurate, then words that resemble the binding profile should score highly in this analysis.

For *hif-1* and *nhr-23*, the most significantly enriched words in the positively and negatively changing genes, respectively, matched the canonical binding sites (Table 2). A word matching the *hlh-30* binding site scored 6^th^ overall among the up-regulated genes, and for *daf-12*, four of the top 20 words for the up-regulated genes contained GAACT or AACTT. These partially match the reverse compliment of a reported *daf-12* binding half-site, AGTTCA^47^. In the *daf-16* data set, several words matching the so-called “*daf-16* associated element” (DAE) scored highly. However, none of the four words matching the canonical *daf-16* binding site, T(G/A)TTTAC, and its reverse compliment were among the words comprising our promoter-word dictionary, precluding these from being represented in the analysis. The canonical binding site for the final transcription factor, *lin-14*, is GAAC, but like the canonical *daf-16* binding site, neither this word nor its reverse compliment was present in the promoter word dictionary, precluding it from representation. Even if GAAC had been present in the dictionary, however, it likely would not have reached statistical significance, as a word comprising only four bases would be expected to occur in most promoters by random chance (see Discussion). Taken together, these results suggest that module weighted regulatory sequences can be used to determine important regulatory sites in a gene expression experiment, and further validate our method for calculating weighted annotations.

### Module-weighted regulatory sequence analysis suggests *hif-1* plays a key role in the effect of *isp-1* mutation

To further test whether module-weighted regulatory sequences could provide biological insights into gene expression data, we used them to analyze a published microarray dataset for *C. elegans* carrying mutations in *isp-1* (iron-sulfur protein, respiratory complex III)^48^. Reduction-of-function *isp-1* mutations extend lifespan in a *hif-1-*dependent fashion^13–15^, but, unexpectedly, we found that the overlap among the significant genes of microarray measurements comparing each mutant to wild type was not statistically significant (*X_2_* test p-value = 0.17; Fig. S4). However, when we projected the full set of gene expression changes in *hif-1* and *isp-1* mutants into gene module space, we found that these projections were strongly anti-correlated (Pearson correlation between SVE = −0.730, p = 4.7E-36). (The *hif-1* and *isp-1* gene expression data was not used in our module construction, as it was not on the Affimetrix platform.)

To determine how often two sets of gene fold changes generated by different experiments could be expected to show this degree of correlation when projected into gene module space, we determined projection correlations for all possible pairs of the 716 Affymetrix contrasts (described above), excluding pairs in which both contrasts originated from the same experiment (i.e., from the same GEO series). This produced 188,805 contrast pairs, 13,376 (7.08%) of which showed a statistically significant correlation (Holm corrected p-value < 0.05). The strength of the *hif-1* and *isp-1* correlation would rank 1570^th^, i.e., within the top 1%, had those experiments been part of the (Affymetrix only) contrast set. The full set of contrast comparisons are provided as Supplemental Data, as contrasts that are similar in gene module space but generated by different experiments could prove useful to others for hypothesis generation (e.g., had we observed the similarity between the *hif-1* and *isp-1* projections, we might have hypothesized a role for *hif-1* in *isp-1* mutants before such a role was suspected.)

We next calculated promoter word z-scores for the *isp-1* microarray data set using our method and compared the results to those for *hif-1.* We observed a very strong correlation between the promoter word z-scores for *isp-1* mutants and those for *hif-1* mutants (Fig. 4b, R = −0.581, p < 2.2E-16). The correlation was negative, consistent with the interpretation that the life extension observed when *isp-1* activity is reduced requires activation of gene expression by HIF-1. The strong promoter word z-score correlation between these datasets, and the coincidence of their projections in our modules despite a lack of similarity among their most differentially expressed genes, suggests that the role of HIF-1 in regulating the lifespan of *isp-1* mutants may be to instigate small but coordinated expression changes in many genes, most of which fail significance tests for differential expression in one or both datasets. In general, it would be interesting to learn to what extent this situation, which would not be detected by many genetic or bioinformatic methods, has arisen during the evolution of gene circuits.

## Discussion

Scientists that study biology at the sub-cellular level face the challenge of understanding a world that operates on a different time and size scale than the one we perceive in our everyday experience. It is a world that has mainly been observed indirectly and in isolated fragments, and the descriptions that have been made of it likely fail to capture parts of its essence. For this and other reasons, some advocate the development of mathematically driven descriptions of genes and gene products based on data rich assays, such as computationally predicted gene modules. These can aid the interpretability of gene expression assays by reducing the dimensionality of a data set from thousands of significant genes to perhaps tens of significant modules. Of course, this leaves the researcher with the challenge of interpreting gene module activities, e.g. by checking for enriched annotations among significant modules’ genes.

Alternatively, as we have shown here, gene module predictions can be combined with prior knowledge (Boolean annotations) to generate module-weighted annotations. Analyses performed with module-weighted annotations leverage the power of gene modules but provide the researcher with results of a more familiar form, e.g., significant GO categories. Such analyses may prove more informative than those based on Boolean annotations in many cases. In our comparison of module-weighted annotation analysis to GSEA, experiments that were highly similar at the gene level bore greater similarity in annotation significance levels with our method than with GSEA, and highly dissimilar experiments bore less. These results suggest that module-weighted annotation analysis may be less prone to the confounding effect of gene-gene correlations and serve to motivate its further development, work that is currently ongoing by our lab.

Whereas ICA has been applied to the prediction of gene modules before, we could find no examples in the literature of optimizing the number of extracted components in the manner that we describe. Combined with the improved ability to partition independent components provided by our artificial neural network approach, we expect that our results will spur exploration into additional applications of ICA to the analysis of biological data. In addition, we expect the regulatory word analysis we describe to stimulate many specific, testable hypotheses about the roles of specific transcription factors in biological processes. For example, had we not known previously that HIF-1 regulated life extension in *isp-1* mutants, we would have generated this hypothesis upon observing the strong similarity between promoter word scores in the *isp-1* and *hif-1* microarray datasets. Thus this method serves as a discovery tool that can link together seemingly disparate factors (such as *hif-1* and *isp-1),* or even prompt new searches for functional significance when gene modules that cannot described satisfactorily by existing annotation terms.

Accurate estimates of module-weighted annotations depend on accurate estimates of gene modules. While we have taken steps to maximize module prediction accuracy for the microarray compendium we assembled, many additional gene modules may exist in ***C. elegans*** that were not perturbed sufficiently in the samples comprising the compendium to be detected. These gene modules would remain hidden, and would therefore fail to exert an effect on annotation weights. As new areas of research are explored and new experiments are published, however, new gene modules may be discovered and module-weighted annotations can be recalculated to improve estimated weights. We envision a system in which this would occur automatically as new experimental data is deposited into the GEO database.

Four of the six transcription factor perturbation microarray experiments we analyzed produced high scoring promoter words that closely matched the known DNA binding sites of the corresponding transcription factors. For the two that did not, *daf-16* and *lin-14*, words that exactly matched the factors’ canonical binding sites were not present in the promoter word dictionary generated by the Mobydick algorithm, and results for these factors could be poor for this reason. Another possible explanation, however, stems from our method for calculating word enrichment among module promoters. We calculated the p-value for module-wise enrichment based on the presence or absence of each word in each gene’s promoter. Thus, words that are present many times in a gene’s promoter do not contribute anything more to the p-value calculation than words that are present only once. The canonical DAF-16 binding site occurs in approximately 50% of all 2k-bp gene promoters, but in the 12 genes with the largest expression changes in *daf-2* mutants, in which DAF-16 becomes activated, Zhang et al.^49^, found that the mean number of occurrences of the DAF-16 binding site is 5.1. Thus, a single copy of the DAF-16 binding site may be insufficient to confer regulation by this factor. A modification of our method that uses gene-wise promoter word enrichment rather than presence vs. absence may prove to be more accurate for predicting active regulatory sequences for transcription factors with highly abundant binding sites, such as DAF-16; this is a question we plan to address in future work. Interestingly, the second highest scoring word for the genes down-regulated upon *daf-16* perturbation was GGAAG, and this word occurs twice more as a substring among the top 20 scoring words. This sequence is a partial match to an alternative *daf-16* binding site reported in hookworm^50^, G(A/G)(C/G)A(A/T)G, suggesting that this site may be functional in *C. elegans* as well.

In addition to its utility in analyzing gene expression data, the module-weighted regulatory sequence matrix could be used to identify transcription factor target genes, providing that words could successfully be matched to transcription factors. Alternatively, ICA of this matrix could be used to find words that “travel together” in terms of their predicted influence on genes. Such independent components, comprising weighted words rather than weighted genes, would presumably represent a set of possible binding sites for a factor, similar to a sequence logo or position-specific weight matrix. Preliminary work on generating word independent components showed that palindrome pairs often scored highly together, lending credence to this approach. Given the dearth of well-characterized transcription factor binding sites in *C. elegans,* matching word-independent components to transcription factors still presents a challenge. We expect the wealth of transcription factor binding data currently being generated by the modENCODE consortium and others, however, to be extremely useful toward this end.

## Methods

### Compendium construction

To build our compendium of 1386 *C. elegans* Affymetrix arrays, we first downloaded all CEL files with the appropriate platform ID (GPL200) from the GEO database available on March 1, 2014, excluding those for which the organism was not *C. elegans* and the sample type was not RNA. We excluded arrays from experiments for which fewer than 8 hybridizations were performed in order to mitigate the effect that under-sampled conditions might have on predicted modules. We then performed a quality control step using the quality assessment functions provided in the *simpleAffy* (v2.40.0) R package (http://bioinformatics.picr.man.ac.uk/simpleaffy/), discarding arrays that did not meet the quality thresholds recommended in the *simpleAffy* documentation.

We generated expression values for probesets separately for each experiment (determined by GEO series IDs) using the RMA preprocessing procedure provided in the *affy* (v1.40.0) R package ^51^, then used the *bias* (v0.0.5) R package^52^ to remove intensity-dependent biases in expression levels. We then concatenated the expression matrices for each experiment into a single matrix. Next, we either performed between-experiment quantile normalization^53^ on the entire matrix using the *limma* (v3.18.13) R package^54^, or omitted this step, depending on preprocessing method to be tested. Finally, we scaled and centered the arrays and centered the genes such that the mean of each row and column were zero and the standard deviation of each array was 1.

### Conducting ICA

To conduct ICA of the gene expression matrix, we used the ***fastICA*** (v1.2-0) R package (http://CRAN.R-project.org/package=fastICA) with default parameters except for the “method” parameter, which we set to “C” to increase computational speed, and the “row.norm” parameter, which we set to “TRUE” in order to balance the total compendium variance between genes with subtle changes in expression values and those with large changes in expression values. We used the same parameters to conduct ICA of the word / module p-value matrix.

### Partitioning of independent components

To convert independent components to discrete sets of genes, we employed two methods. In the first, for each component, we assigned all genes with a weight <= −3 to the negative hemi-module, and all genes with a weight >= 3 to the positive hemi-module. In the second, we created an artificial neural network using the *neuralnet* (v1.32) R package (http://CRAN.R-project.org/package=neuralnet) to predict positive and negative partitioning thresholds for each independent component, based on the component’s skewness and kurtosis (see Supplemental Methods), then assigned genes whose weights exceeded these thresholds to the corresponding hemi-modules.

### Obtaining gene annotations and additional microarray data

To obtain GO term and REACTOME pathway annotations for genes we used the *biomaRt* (v2.18.0) R package^55,56^, using the *ensembl* mart for data retrieval. To obtain tissue annotations for *C. elegans* genes, we downloaded all available data from the GFP Worm database (http://gfpweb.aecom.yu.edu/)^57^, which contains annotated expression patterns of promoter::GFP fusion constructs; in total, this dataset provided annotations for 1821 genes across 89 tissue types ***(n.b.,*** we considered the same tissue in different development stages to be distinct tissue types). To obtain expression data from a different platform for use in optimization of gene module prediction, we downloaded the fold change matrices for all GEO series conducted on the Washington University *C. elegans* 22k 60-mer array (GEO platform ID: GPL4038), a two-color spotted array platform, and concatenated these column-wise into a single matrix. To obtain microarray data for *nhr-23(RNAi),* we downloaded gene fold changes for the GEO series GSE32031, which contains three control samples and three *nhr-23(RNAi)* samples^58^; gene fold changes were calculated using the GEO2R web service (http://www.ncbi.nlm.nih.gov/geo/geo2r/). To obtain fold changes for *isp-1* mutants, we used data previously published by our group in which *isp-1(qm150)* mutants were compared to wild type controls^48^. To obtain fold changes for *hif-1* mutants, we used the *maanova* (v1.33.2) R package (http://research.jax.org/faculty/churchill) and data previously published by Shen, et al.^46^, to calculate the induced gene fold changes upon mutation of *hif-1.*

### Optimizing gene module prediction

To optimize gene module prediction, we performed ICA with different parameters, varying the number of extracted components from 5 to 500 by increments of 5 and varying the compendium between one generated with between-experiment quantile normalization and one generated without this step. For each parameter combination, we repeated ICA 5 times, for a total of 1000 ICA runs.

We tested the biological validity of the independent components generated by each ICA run by determining the number of annotations that were enriched in at least one hemi-module. To make this determination, we first calculated a p-value for the enrichment of genes associated with each annotation term in each hemi-module using the hypergeometric test. We then applied the Simes method^40^ for multiple hypothesis testing (alpha = 0.05) to the set of p-values for each annotation term; failure of this test indicates that at least one of the p-values is truly significant. To verify the accuracy of our module quality statistics, we repeated all tests using module definition matrices in which gene IDs had been randomly shuffled.

To test the ability of a set of independent components to represent data from a different microarray platform, we first projected the data from the second platform onto the independent components (see below). This operation produces a mixing matrix, which may be interpreted as describing the weight of each independent component in each of the projected microarrays. We then attempted to recover the original data by determining the dot product of the module definition matrix and the mixing matrix. We compared this matrix with the original matrix and calculated the root mean squared deviation (RMSD) between the two. We normalized this value by dividing by the range of values between the two matrices, resulting in NRMSD.

### Statistical testing of module 3’-UTR length bias

We observed that ***C. elegans*** 3’-UTR lengths are approximately log-normally distributed (Figure S5). Therefore, we chose to use the log of each 3’-UTR length in our calculations to allow the use of parametric statistical tests, such as Student’s t-test. For those genes with multiple annotated 3’-UTRs, we determined the log of the individual 3’-UTR lengths and used the mean of these numbers for the gene’s 3’-UTR length.

In our statistical test for 3’-UTR length biases in predicted modules, we first calculated the weighted mean ***C. elegans*** 3’-UTR length. We weighted each gene’s contribution to this mean by the frequency with which it appears in our predicted modules in order to adjust for different propensities for module inclusion by different genes. We then used one-sample t-tests to calculate p-values for whether the mean 3’-UTR length of each hemi-module differs significantly from the weighted mean ***C. elegans*** 3’-UTR length. We used the Benjamini-Hochberg procedure on these p-values to control the false discovery rate at a level of 0.1.

### Generation of *X_a_* matrix (annotation / hemi-module enrichment score matrix)

To generate the *X_a_* matrix, we first created gene sets from the module definition matrix, ***S_g_***, using ANN-based partitioning. This produced two gene sets (which we refer to as hemi-modules) per gene module, for a total of 418. We then calculated a hypergeometric probability for each word in each hemi-module, using the frequency of genes bearing a particular word in their promoter in the hemi-module, the frequency of such genes in the compendium, the number of genes in the hemi-module, and the number of genes not in the hemi-module as the *q, m, k,* and *n* input parameters, respectively, to the *phyper()* function of the *stats* (v3.0.3) R package (http://www.R-project.org/).

We used these p-values to populate a matrix with a column for each hemi-module and a row for each word in our promoter dictionary. For under-represented words, we entered the *log(p-value)* in the matrix, and for over-represented words we entered the *–log(p-value).*

### Weighted annotation enrichment analysis

To test whether a set of gene fold-changes were significantly enriched for specific annotations, given a module-weighted annotation matrix, ***X_w_,*** we calculated the dot product of the data vector, ***x***, comprising the set of gene fold-changes, and the ***X_w_*** matrix:

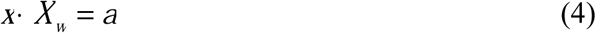

The resulting vector, ***a***, provides an indication of the degree to which genes with strong weights for each annotation also have strong fold-changes. To generate p-values from these, we permuted the fold-change vector, ***x***, 1000 times to create a background distribution for each annotation, which we then used to determine z-scores.

### Data projection and calculation of SVE

To project a data vector, ***x***, such as a set of gene expression fold changes, onto a set of gene modules (or a weighted annotation matrix), we used the scalar projection method, in which a mixing vector, ***a***, is calculated from the dot product of the data vector and the transpose of the unit vectors comprising the module definitions, *Ŝ*^T^, as shown in equation 5:

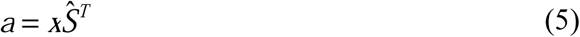

The resulting mixing vector, ***a***, provides an indication of the weight of each module definition vector in the projected data, ***x***. Projection of a data matrix, ***X***, which generates a mixing matrix, ***A***, was carried out using the same procedure.

To calculate signed variance explained (SVE), we calculated the relative variance explained (VE) for each module from ***a*** as follows:

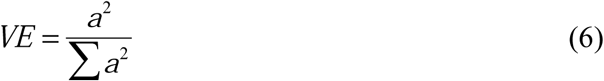

We then multiplied these values, which are strictly positive, by −1 in each case where ***a*** < 0 to obtain SVE.

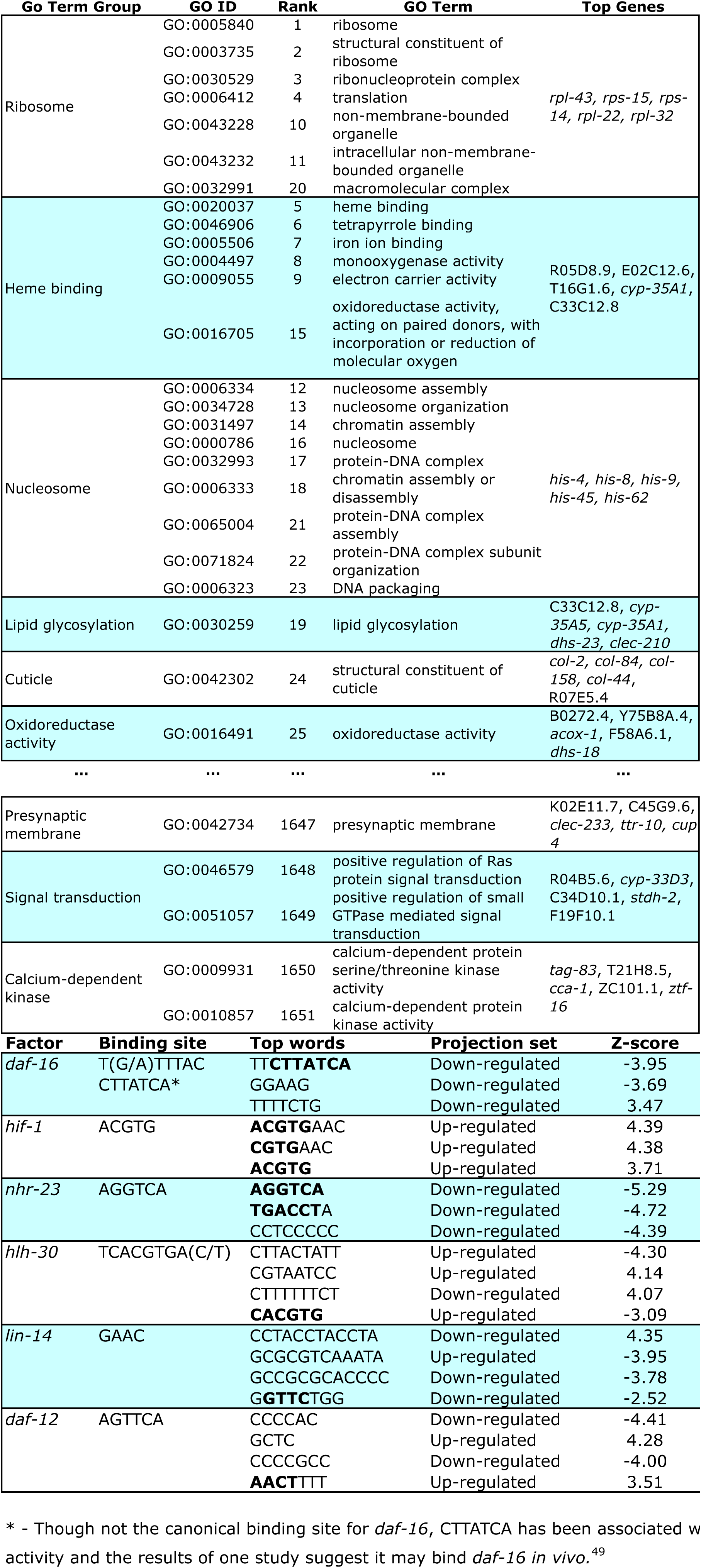

## Supplemental methods

### Generation of simulated data

To generate a simulated microarray compendium containing a proscribed number, *n*, of true gene modules, we performed ICA on a test compendium and extracted 2*n* independent components. We then randomly selected *n* rows of the resulting S matrix and multiplied these with the corresponding columns of the A matrix. This generated a matrix with the same dimensions as the test compendium, but with only *n* latent gene modules. We then added random Gaussian noise to each data point in this matrix. Increasing the level of noise decreased the accuracy of modules extracted from the matrix, but did not alter the observed property that accuracy reached a maximum when the number of extracted components matched the number of latent modules (data not shown).

### Construction of Mobydick dictionaries

To construct promoter and 3’-UTR dictionaries, we ran the Mobydick^39^ program once on the complete set of *C. elegans* promoters, using DNA sequence from the transcription start site to 2000 b.p. upstream for each gene, and again on the complete set of 3’-UTRs with lengths of at least 25 n.t. Sequences were obtained using the *biomaRt* (ver 2.14.0) R package^55,56^. Application of Mobydick to promoter sequences produced a dictionary of 5230 words, and application to 3’-UTR sequences produced a dictionary of 968 words.

### Calculation of significance of 3’-UTR word enrichment

Because 3’-UTRs differ in length, and because gene modules show a tendency toward inclusion of genes with similar length 3’-UTRs, calculation of the enrichment of 3’-UTR words in module genes required a length-normalization step. To achieve this, we applied the method described van Helden, et al.^59^ Briefly, we determined the per nucleotide frequency of each word in the entire set of 3’-UTRs, then used the binomial distribution to determine whether each word occurs more often than expected by random chance in a sequence, given the number of occurrences of the word in the sequence and the sequence length. We then applied the Holm-Bonferroni correction to the resulting p-values and marked all words with a corrected p-value < 0.5 as present in the 3’-UTR.

### Generation of Affymetrix contrasts

To generate the 716 Affymetrix contrasts, we examined all *C. elegans* Affymetrix experiments (organized as GEO series) available for download in the GEO database on January 23, 2015. For each experiment, we generated contrasts (two sets of microarrays to compare to each other) based on the following rules: 1) we compared genetic mutants to wild type controls, or to the genotype most resembling wild type if true wild type animals were not used in the experiment, 2) we compared animals subjected to a treatment, such as an environmental stress, to animals with the same genotype but subjected to control conditions, 3) we compared animals harvested at different time points to animals with the same genotype but harvested at the earliest time point represented in the experiment. When necessary, we divided large experiments into smaller subsets in order to simplify contrast generation and facilitate the application of the above rules. We generated fold-changes from contrasts using the RMA procedure provided in the *affy* (v1.40.0) R package.

### Generation of artificial neural network for independent component partitioning

To create an artificial neural network for use in partitioning independent components, we first generated simulated data to use as test, training, and validation sets. We generated this data by first randomly permuting the expression values of 100 arrays comprising our *C. elegans* microarray compendium column-wise to create a background devoid of non-random signal, but with a similar gene expression value distribution to real data.

Into this random background we inserted simulated gene modules by first picking a gene to use as a module seed pattern, then changing the expression values of other genes such that they positively or negatively correlated with the expression values of this gene across all or a subset of arrays. We varied the number of genes comprising the simulated module, the strength of adherence of each gene to the seed pattern, the fraction of genes within the module with positive and negative correlation to the seed pattern, and the number of arrays in which this correlation existed. In all, we generated over 10,000 random modules and inserted them into separate sets of random background arrays, so that each array set would contain a single non-random module.

We then attempted to recover each simulated module using ICA. We extracted a single component from each simulated array set and deemed the extraction successful if 3 of the top 5 most strongly weighted genes in this component were in the simulated module. For successful extractions, we calculated the optimal partitioning thresholds for the positive and negative hemi-modules, as well as the skewness and kurtosis of the module definition vector using the *moments* (v0.13) R package (http://CRAN.R-project.org/package=moments).

Using this data, we trained an artificial network to predict the optimal partitioning thresholds for an independent component from the skewness and kurtosis of its gene weights using the *neuralnet* (v1.32) R package (http://CRAN.R-project.org/package=neuralnet). We generated another simulated module set in the same manner as the first to use as a test set, and varied the architecture of the artificial neural network until the prediction performance reached a maximum value. This occurred when the artificial neural network contained two hidden layers, each with 11 nodes. We confirmed that the artificial neural network was not over-fit to the test set by measuring its performance in a third set of simulated data, the validation set. Performance on this set was similar to that on the test set. The structure of this artificial neural network is shown in Figure S6; an R data file containing the artificial neural network is available for download on our website (http://kenyonlab.ucsf.edu/data/ann.11.11.Rdata).

## Acknowledgements

We would like to thank H. Li, H. El-Samad, and H. Madhani for critical review and discussion of this work. This work was supported by the NIH R36/R01 grant AG011816 to C. K.

## Competing interests statement

The authors declare that they have no competing financial interests.

